# Acidophilic Micrarchaeon Seems to Maintain a Slightly Alkaline Cytosolic pH

**DOI:** 10.1101/2021.05.02.442328

**Authors:** Dennis Winkler, Sabrina Gfrerer, Johannes Gescher

## Abstract

Despite several discoveries in recent years, the physiology of acidophilic Micrarchaeota remains largely enigmatic. “*Candidatus* Micrarchaeum harzensis A_DKE”, for example, highly expresses numerous genes encoding hypothetical proteins and their function is difficult to elucidate due to a lacking genetic system. Still, not even the intracellular pH value of A_DKE is known, and heterologous production attempts are generally missing so far. Hence, A_DKE’s isocitrate dehydrogenase (*Mh*IDH) was recombinantly produced in *Escherichia coli* and purified for bio-chemical characterisation. MhIDH appeared to be specific for NADP^+^, yet promiscuous regarding divalent cations as cofactors. Kinetic studies showed *K*_*M*_-values of 53.03±5.63 µM and 1.94±0.12 mM and *k*_*cat*_-values of 38.48±1.62 s^-1^ and 43.99±1.46 s^-1^ for DL-isocitrate and NADP^+^, respectively. *Mh*IDH’s exceptionally low affinity for NADP^+^, potentially limiting its reaction rate, can be likely attributed to the presence of a proline residue in the NADP^+^ binding-pocket, which might cause a decrease in hydrogen bonding of the cofactor and a distortion of local secondary structure. Furthermore, a pH optimum of 7.89 implies, that A_DKE applies potent mechanisms of proton homoeostasis, to maintain a slightly alkaline cytosolic milieu in a highly acidic environment.

## 1. Introduction

Microorganisms can survive and thrive under extreme environmental conditions [1–3]. Bacteria and Archaea in particular are often adapted to niches of extreme temperature, pressure, radiation, salinity or pH, which allows them to populate a vast variety of habitats inaccessible to non-extremophiles [1,4]. Still, to cope with these conditions, requires a significant amount of metabolic resources, in order to adjust the intracellular reaction conditions.

Acidophilic Archaea, for example, are thriving in environments with pH values below pH 3 [5,6], in extreme cases, optimal growth occurs close to pH 0 (i.e. *Picrophilus torridus* [7], *Ferroplasma acidiphilum* [8]). Still, these organisms are able to maintain less acidic to near neutral internal pH (pH_i_) values between pH 4.6 (i.e. *Picrophilus torridus*, [9]) and pH 6.5 (i.e. *Sulfolobus acidocaldarius*, [10]) by applying numerous synergistic mechanisms of proton homoeostasis [5,11,12].

Micrarchaeota were originally discovered in habitats with pH values between 0.5 and 4.0 [13]. Common characteristics of known members of this phylum are small-sized, circular genomes and an overall limited metabolic potential [13–17]. Thus, Micrarchaeota are assumed to be dependent on a symbiotic relationship with host organisms of the order *Thermoplasmatales* [15,18,19].

To our best knowledge, the only acidophilic Micrarchaeon currently cultivated under laboratory conditions is “*Candidatus* Micrarchaeum harzensis A_DKE” in co-culture with its putative host “*Ca*. Scheffleriplasma hospitalis B_DKE” [17]. The culture was enriched from acid mine drainage biofilms originating from the abandoned pyrite mine “Drei Kronen und Ehrt” in the Harz Mountains (Germany) [18,20]. Optimal growth of the laboratory culture was achieved at pH 2 [17]. Although an extensive multi-omic-approach, comprising genomics, transcriptomics, proteomics, and metabolomics, has been conducted on A_DKE [17,21], details of its metabolism still remain enigmatic. Approximately a third of the genes in the A_DKE genome encode hypothetical proteins, most of which are also actively expressed, according to transcriptomic data [17]. Of note, these hypothetical protein-encoding genes comprise 35 % and 60 % of A_DKE’s 100 and 10 highest expressed genes, respectively [unpublished data]. Considering A_DKEs reduced genome and so far largely enigmatic metabolism [17,18], these proteins of unknown function might be crucial for understanding A_DKE’s physiology. Yet, due to low sequence conservation, *in silico* characterisation of these proteins is currently not possible and thus biochemical characterisation remains key to fully understand A_DKE’s physiology. Investigating the function of these proteins by means of heterologous expression proves to be difficult, since there is no information on the intracellular conditions in Micrarchaeota. One of the factors defining the intracellular conditions and protein stability of an organism is the pH_i_ value, as it affects the activity of proteins, for example in DNA transcription, protein synthesis and biocatalysis (for reviews, please check [11,22]). Thus, a suitable production platform must be chosen mimicking the intracellular conditions of A_DKE as best as possible to facilitate proper folding of the proteins of interest.

The goal of this study was to gain evidence for the pH_i_ of A_DKE by biochemical characterisation of an intracellular enzyme. As a target protein, its isocitrate dehydrogenase (IDH) was chosen, which is a key enzyme of the tricarboxylic acid cycle catalysing the oxidative decarboxylation of isocitrate to α-ketoglutarate and CO_2_ [23]. This analysis revealed a slightly alkaline pH optimum indicating that A_DKE displays a comparatively high pH_i_ for an acidophile.

## 2. Materials and Methods

### 2.1 Database Research and Bioinformatic Sequence and Structure Analyses

Genomic (accession number: CP060530) and transcriptomic data (accession numbers: SRX8933312-SRX8933315) of A_DKE were accessed via the National Center for Biotechnology Information NCBI [24] (bio project number: PRJNA639692). The pH optima and kinetic parameters of homologous enzymes for comparison with experimentally identified parameters for *Mh*IDH were obtained from the BRENDA database ([25], www.brenda-enzymes.org).

The theoretical molecular weight and isoelectric point of *Mh*IDH were calculated using the CLC Main Workbench 20.0.1 (QIAGEN, Aarhus, Denmark). Conserved sequence motifs and protein domains were detected using the Pfam database ([26], www.pfam.xfam.org). *Mh*IDH homologues were identified via BLASTp [27] search of the UniprotKB/swiss-prot database [24] via NCBI. A multiple sequence alignment comparing *Mh*IDH with experimentally verified homologues from *Escherichia coli* K-12 (*Ec*IDH, NCBI: P08200.1), *Aeropyrum pernix* K1 (*Ap*IDH, NCBI: GBF08417.1), *Archaeoglobus fulgidus* DSM 4304 (*Af*IDH, NCBI: O29610.1), *Haloferax volcanii* DS2 (*Hv*IDH, NCBI: D4GU92.1) and *Sulfolobus tokodaii* Strain 7 (*St*IDH, NCBI: BAB67271.1) was carried out using the Clustal Omega algorithm [28–30] as a plugin for the CLC Main Workbench 20.0.1. The alignment was visualised using the ESPript 3.0 server ([31], www.espript.ibcp.fr).

Homology modelling of a putative *Mh*IDH structure was achieved via the CLC Main Workbench 20.0.1 using the crystal structure of *Ec*IDH in complex with Ca^2+^, isocitric acid and NADP^+^ ([32], PDB: 4AJ3, 49.5 % homology, 1.9 Å resolution) as a template. Assessment of local model quality and B-factor, as well as docking of the cofactors Mn^2+^, NADP^+^ and the substrate isocitrate to the *Mh*IDH model structure was performed using the ResQ server [33] and the COACH server [34,35] respectively. Protein ligand interactions were analysed using the PLIP server ([36], www.plip-tool.biotec.tu-dresden.de/plip-web). All protein structures were visualised using PyMOL 2.3.3 (Schrödinger, Ney York, USA).

### 2.2 Cloning and Recombinant Expression of *icd*2_6*x His*_

The *icd2* gene was PCR-amplified from genomic DNA isolated from a co-culture containing “*Ca*. Micrarchaeum harzensis A_DKE” and “*Ca*. Scheffleriplasma hospitalis B_DKE” [17] via oligonucleotide primers 1 & 2 (see Table 1). The latter introduced a 6x His-tag encoding sequence to the 5’-end, as well as complementary overlaps to the target vector pBAD202 (Invitrogen, Carlsbad, CA, USA). pBAD202 was linearised via inverse PCR using primers 3 & 4 (see Table 1). Both PCR products were gel-purified using the Wizard® SV Gel and PCR Clean-Up System (Promega, Mannheim, Germany) and assembled via isothermal in vitro ligation [37]. The resulting plasmid pBAD202_*icd2*_*6x His*_ was transformed into *E. coli* Rosetta pRARE (Merck, Darmstadt, Germany).

**Table 1.**
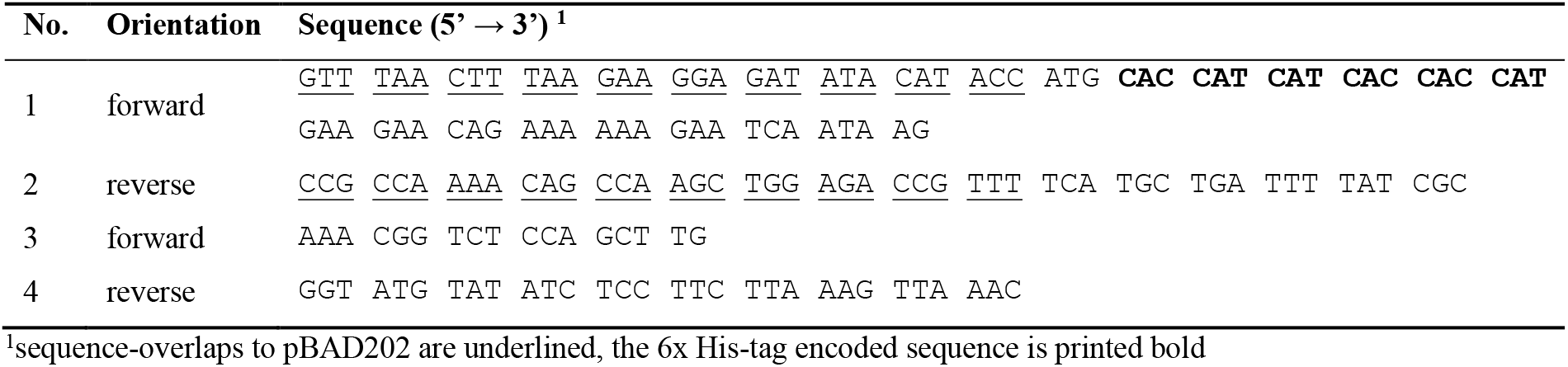
Oligonucleotide primers used in this study.

In order to monitor production of *Mh*IDH_6x His_ over time, *E. coli* Rosetta pRARE pBAD202_*icd2*_*6x His*_ was cultivated in shaking flasks containing 50 mL Terrific Broth medium (1.2 % (w/v) tryptone, 2.4 % (w/v) yeast extract, 0.5 % (w/v) glycerol, 17 mM KH2PO4, 72 mM K2HPO4) supplemented with 50 µg mL^-1^ kanamycin and 30 µg mL^-1^ chloramphenicol at 37 °C and 180 rpm. Upon reaching an OD_600_ of 0.6-0.8, expression of *icd2*_*6x His*_ was induced by addition of 1 mM L-(+)-arabinose. From this point forth, the culture was incubated at 30 °C and 180 rpm and samples (1 mL) were taken at different time points after induction (0, 1, 2, 4, 6 and 24 h), and subjected to OD_600_-measurement using a GENESYSTM 20 spectrophotometer (Thermo Fisher Scientific, Schwerte, Germany) and preparation for SDS-PAGE analysis. Samples were centrifuged for 2 min at 16 000 g and cell pellets were resuspended in 75 µL of 2x SDS loading dye (240 mM TRIS/HCl (pH 6.8), 20 % (v/v) glycerol, 2 % (w/v) SDS, 100 mM DTT, 0.02 % (w/v) Orange G) per OD600 of 0.2, boiled for 10 min at 95 °C and centrifuged for 5 min at 16 000 g. After determination of the optimal induction time, over-expression was carried out in a total volume of 1 L as described above. Cells were harvested for 15 min at 16 000 g and 4 °C, 4 h after induction and stored at 20 °C until used.

### 2.3 Isolation and Affinity Purification of *Mh*IDH_6*x His*_

The cell pellet of an expression culture was resuspended in IMAC buffer (50 mM HEPES/NaOH (pH 7.4), 500 mM NaCl) followed by the addition of a spatula tip of Deoxyribonuclease I (SERVA Electrophoresis, Heidelberg, Germany). Cell extracts were prepared using mechanical disruption in an FA-078 FRENCH® Pressure Cell Press (SLM Aminco, Urbana, IL, USA) at 137.8 MPa. The raw lysate was fractioned by suc-cessive steps of centrifugation. Intact cells and cell debris were pelleted for 15 min at 6 000 g and 4 °C. Membranes were separated from the plasma fraction via ultracentrifugation for 60 min at 138 000 g and 4 °C. The membrane pellet was resuspended in solubilisation buffer (20 mM HEPES/NaOH (pH 8.0), 150 mM NaCl, 2 % (v/v) Triton X-100) and the plasma fraction was passed through a 0.2 µm syringe filter (Sarstedt, Nümbrecht, Germany) to remove remaining insoluble particles. Samples of the raw lysate, as well as the membrane and plasma fraction were used for SDS-PAGE.

Nickel Immobilised Metal Ion Affinity chromatography (Ni^2+^-IMAC) for protein purification was conducted using a HisTrap® HP 5 mL column (GE Healthcare, Munich, Germany) coupled to a BioLogic DuoFlow™ Chromatography System (Bio-Rad, Munich, Germany). The column was equilibrated with IMAC buffer, prior to loading with plasma fraction. Non-specifically bound proteins were removed by washing with IMAC buffer containing 80 mM imidazole. Elution of the target protein was achieved with IMAC buffer containing 500 mM imidazole. The eluted fraction was concentrated using a 3 kDa MWCO centrifugal filter (Merck, Darmstadt, Germany). Samples of the column flow-through, wash and eluate were used for SDS-PAGE.

Size exclusion chromatography (SEC) of the concentrated protein solution was conducted using a HiLoad™ 26/600 Superdex™ 200 pg column (GE Healthcare, Mu-nich, Germany) coupled to the aforementioned chromatography system. The column was equilibrated and run isocratically with IDH buffer (50 mM HEPES/NaOH (pH 7.4), 150 mM NaCl, 1 mM DTT, 0.5 mM MgCl_2_). The eluted fractions were collected, concentrated and analysed via SDS-PAGE. For long term storage at -20 °C, 50 % (v/v) glycerol was added.

### 2.4 Protein Quantification, SDS-PAGE & Western Blot

Protein quantification of samples collected for analysis via SDS-PAGE was carried out according to [38]. Alternatively, purified protein was quantified spectrophotometrically using a NanoDrop 2000 (Thermo Fisher Scientific, Schwerte, Germany).

Samples containing 20 µg of total protein (5 µg in case of purified protein) were mixed with 2x SDS loading dye and separated via denaturing SDS-PAGE in hand cast 12 % TRIS/Glycine gels according to [39]. As reference either BlueStar™ Prestained Protein Marker (NIPPON Genetics, Düren, Germany) or PageRuler™ Prestained Protein Ladder (Thermo Fisher Scientific, Schwerte, Germany) was used. After separation, the gels were subjected to either colloidal staining using Quick Coomassie Stain (Protein Ark, Sheffield, UK) or transfer of the separated proteins to a nitrocellulose membrane (Roth, Karlsruhe, Germany) via a semi-dry blot. The latter was carried out with a Trans-Blot® Turbo™ device (Bio-Rad, Munich, Germany) at 1.3 A for 10 min using a continuous blotting buffer system (330 mM TRIS, 267 mM glycine, 15 % (v/v) ethanol, 5 % (v/v) methanol, pH 8.8).

Densitometric estimation of protein purity from Coomassie-stained acrylamide gels was carried out using the Image Studio Lite 5.2 software (LI-COR, Lincoln, NE, USA).

For immuno-staining the membrane was blocked for at least 1 h at room temperature with TBST (20 mM TRIS/HCl (pH 7.5), 500 mM NaCl, 0.05 % (v/v) Tween® 20) containing 3 % (w/v) skim milk powder. After a few brief rinses with TBST, the blot was incubated with a mouse anti-His-tag primary antibody (Sigma-Aldrich, Steinheim, Germany), diluted 1:1 000 in TBS (10 mM TRIS/HCl (pH 7.5), 150 mM NaCl) containing 3 % (w/v) BSA for 1 h, followed by washing with TBST (4x 5 min) and incubation with a goat anti-mouse alkaline phosphatase secondary antibody (Sigma-Aldrich, Steinheim, Germany) diluted 1:30 000 in TBST containing 3 % (w/v) skim milk powder for 45 min. After washing with TBST (4x 5 min) and several brief rinses with dH_2_O, protein bands were visualised colorimetrically using the AP conjugate substrate kit (Bio-Rad, Munich, Germany) according to manufacturer’s instructions.

### 2.5 Spectrophotometric IDH-Activity Assays and Determination of Kinetic Properties

*Mh*IDH_6x His_ activity and kinetic properties were determined at least in triplicates at 28 °C by monitoring the formation of NADH or NADPH spectrophotometrically at 340 nm using an NADH or NADPH standard curve for quantification. The standard reaction mixture contained 100 mM TRIS/HCl (pH 8.0), 1 mM DL-Na_3_-isocitrate, 5 mM MgCl_2_, 2 mM Na_2_NADP and 0.6-2.5 µg enzyme in a total volume of 200 µL. Each reaction was started individually by addition of either NADP^+^ or enzyme using a TeInjectTM Dispenser (Tecan, Männedorf, Swiss) followed by measurement of A_340_ each 200 ms for 15-30 s using an Infinite® M 200 PRO plate reader (Tecan, Männedorf, Swiss). Investigation of cofactor-specificity was conducted by measuring specific activity with 20 mM NADP^+^ or NAD^+^ in presence of Mg^2+^ and cation-dependency was determined by measuring specific activity in presence of 5 mM MgCl_2_, MnCl_2_, CaCl_2_, ZnCl_2_, NiCl_2_, CuCl_2_, CoCl_2_ and Na_2_EDTA, respectively, with 2 mM NADP^+^. The pH optimum was determined by measuring specific activity in buffers with varying pH values. A corresponding polynomial fitting curve of 5th order was calculated using Origin Pro 2020. In order to span a range from pH 5 to 9.5, three different buffer systems were applied as described in [40]: 0.1 M CH_3_CO_2_Na/CH_3_CO_2_H (pH 5.0-6.0), 0.1 M Na_2_HPO_4_/NaH_2_PO_4_ (pH 5.5-7.5), and 0.1 M TRIS/HCl (pH 7.0-9.5). Enzyme kinetics were determined by measuring the initial reaction rate at increasing concentrations of NADP^+^ (0-10 mM) and DL-isocitrate (0-500 µM), respectively. *K*_*M*_ and *V*_*max*_ were calcu-lated from a non-linear fit based on the Michaelis-Menten model [41,42] using Origin Pro 2020.

## 3. Results and Discussion

### 3.1 *Mh*IDH Shows Conserved Characteristics of Prokaryotic, NADP-Dependent IDHs

pH_i_ values can be estimated from the pH optima of cytosolic enzymes [43]. The enzyme of choice should be monomeric or homo-oligomeric and preferably allow cost-effective and direct activity measurement. The isocitrate dehydrogenase (IDH) of A_DKE fulfils these requirements.

A_DKE possesses only one gene (*icd2*, Micr_00902) annotated to be encoding a putative NADP-dependent IDH, which is actively expressed, according to available transcriptomic data [17]. *In silico* analyses of its amino acid sequence allowed the calculation of a theoretical molecular weight and isoelectric point (pI) of 45.05 kDa and 5.82, respectively, as well as the discovery of a highly conserved isocitrate/isopropylmalate dehydrogenase domain (Pfam: PF00180.20), almost spanning the entire length of the sequence (Thr23-Leu402). Furthermore, a BLASTp search of the UniprotKB/swiss-prot database revealed high sequence homology to several experimentally proven homo-dimeric, NADP-dependent IDHs with nearly all amino acids reported to be involved in substrate and cofactor binding being conserved (see Figure A1 and Table A1). Hence, this bioinformatic data strongly suggests that this protein is indeed an IDH and thus can be used for biochemical characterisation.

### 3.2. *Mh*IDH_6x His_ Can be Produced in *E. coli*

Since direct purification of native *Mh*IDH from “*Ca*. Micrarchaeum harzensis A_DKE” is not feasible due to only low cell density cultures, the corresponding gene was cloned and over-expressed in *E. coli*. Test-expression over time showed high ex-pression levels with a maximum at 4 h after induction and no significant degradation of the product, even 24 h after induction (see Figure 1a). The protein has an apparent molecular weight of roughly 50 kDa, matching the theoretical molecular weight. It was found to be located in the cytoplasmic fraction and could not be detected in the membrane fraction (see Figure 1b). Affinity purification of *Mh*IDH_6x His_ from the plasma fraction was successful in a single step, providing roughly 90 % of electrophoretic homogeneity (see Figure 1c). SEC was used for further purification.

**Figure 1.**
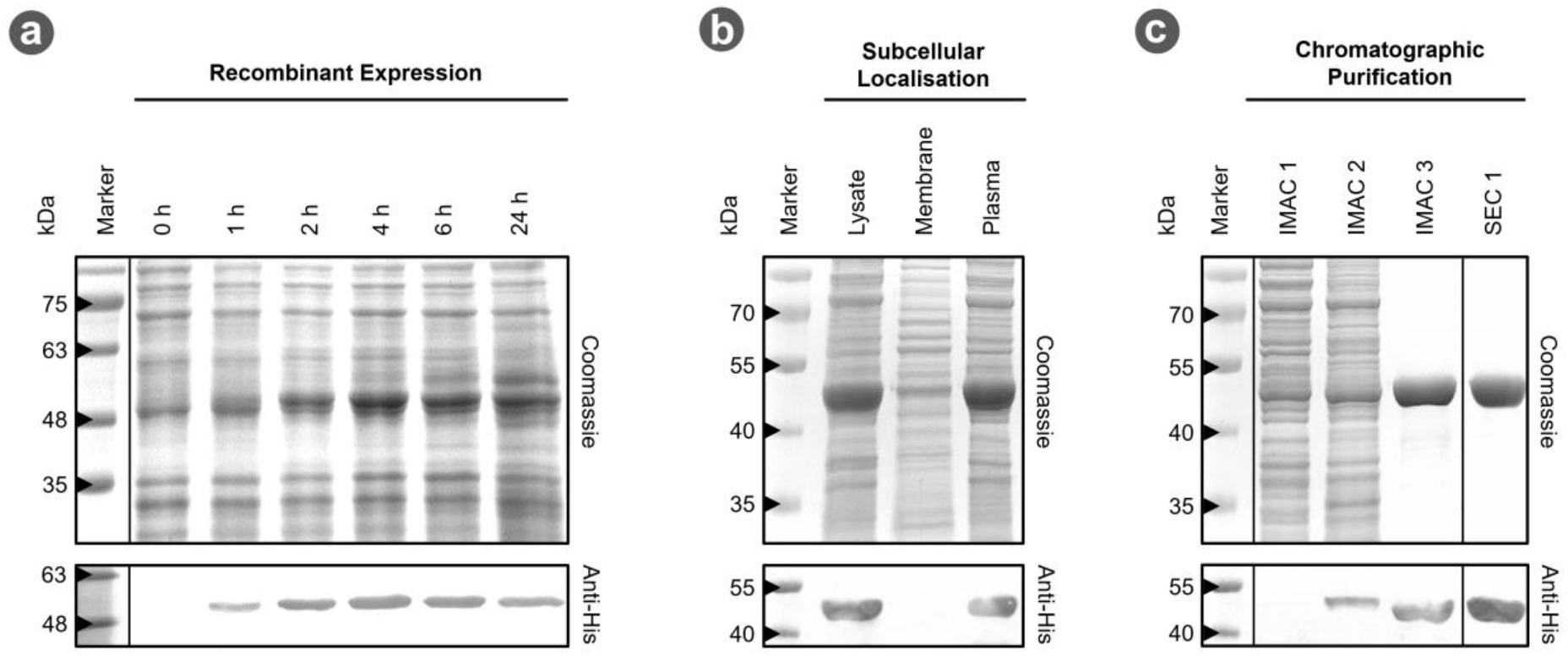
Recombinant production and purification of *Mh*IDH_6x His_. (**a**) 12 % SDS-PAGE of samples from test-expression of *icd2*_*6x His*_. Cell samples were taken 0, 1, 2, 4, 6 and 24 h after induction of gene expression, normalised to identical cell densities and disrupted by thermal and chemical lysis, prior to loading on the gel. Identical gels were prepared for colloidal Coomassie-(top) and colorimetric immuno-staining using an anti His-tag primary antibody (bottom). (**b & c**) 12 % SDS-PAGE of samples from isolation and chromatographic purification of *Mh*IDH_6x His_. Gels were Coomassie- and immuno-stained as described above. IMAC 1, 2, and 3 refer to the flow through during loading of the Ni^2+^-IMAC column, and the fractions which eluted with 80 mM and 500 mM imidazole, respectively. SEC 1 refers to the first fractions eluted during size exclusion chromatography.

### 3.3 Biochemical Properties of *Mh*IDH_6x His_

#### 3.3.1 *Mh*IDH_6x His_ Activity is Dependent on NADP^+^ and Divalent Cations

IDHs catalyse the oxidative decarboxylation of isocitrate to α-ketoglutarate and CO_2_. The electrons released in this process are transferred to either NAD^+^ (EC 1.1.1.41) or NADP^+^ (EC1.1.1.42) [23,44]. Type I IDHs found in Bacteria and Archaea predominantly use NADP^+^ [44,45]. Still, promiscuous forms accepting both cofactors have been reported as well [46–48]. Furthermore, IDHs are known to be dependent on divalent metal cations, such as Mg^2+^ and Mn^2+^ [49]. In order to characterise enzyme activity of recombinant *Mh*IDH, its dependency on different cofactors was tested.

With 41.09±1.02 µmol min^-1^ mg^-1^ *Mh*IDH_6x His_ activity is about 55-fold higher using NADP^+^ as cofactor relative to NAD^+^ with only 0.74±0.09 µmol min^-1^ mg^-1^ (see Figure 2a). The apparent NADP^+^ specificity of the enzyme is also supported by structural data. The primary structure of *Mh*IDH contains conserved amino acid residues (Lys335, Tyr336 and Arg386) in the active site (see Figure A1), which have been shown in *Ec*IDH [50,51], *St*IDH [52] and *Ap*IDH [53] to specifically stabilise the 2’-phosphate moiety of NADP^+^ ensuring that NADP^+^ is bound preferably.

**Figure 2.**
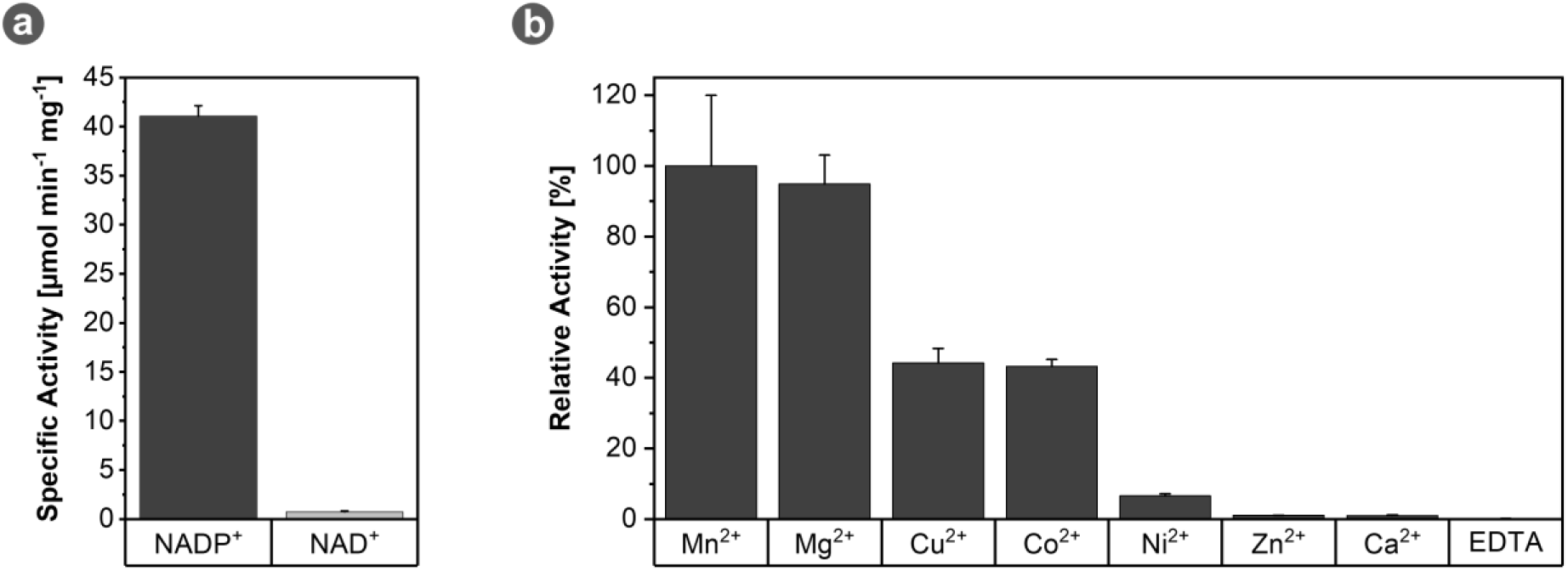
Cofactor-specificity of *Mh*IDH_6x His_. (**a**) Specific IDH activity in presence of 20 mM NADP^+^ (dark grey) or 20 mM NAD^+^ (light grey). Assays were performed at pH 8 and 28 °C in presence of Mg^2+^. (**b**) Relative *Mh*IDH_6x His_ activity in presence of different divalent cations and EDTA. Assays were performed at pH 8 and 28 °C in presence of NADP^+^.

As expected, divalent cations appear to be vital for *Mh*IDH_6x His_ function, as the enzyme does not show any activity in presence of EDTA (see Figure 2b). Still, with several different metal ions having an activating effect, *Mh*IDH is rather promiscuous in this regard. While Mn^2+^ and Mg^2+^ induced maximal activity increases, only 44.2±4.01 %, 43.2±1.99 % and 6.6±0.60 % of relative maximal activity can be achieved with Cu^2+^, Co^2+^, and Ni^2+^, respectively. Zn^2+^ and Ca^2+^, on the other hand, do not seem to enhance enzyme activity, as in presence of these ions *Mh*IDH_6x His_ is only marginally more active than in presence of EDTA. The variance in activation levels in presence of different cations is seemingly independent of ionic radii and is hypothesised to be due to individual modes of binding in the active site of the enzyme [54]. Moreover, Zn^2+^ [55] and Ca^2+^ [54,56] have been reported to inhibit IDH activity. In case of Ca^2+^, this is most likely due to a spatial shift of ligands bound in the active site in order to accommodate the large ionic radius of the cation [56].

#### 3.3.2 *Mh*IDH_6x His_ Shows Highest Activity at Slightly Alkaline pH

With the optimal cofactor combination known, specific activity was measured at different pH values in increments of 0.5. From this data a non-linear fitting curve was calculated with the global maximum of the curve indicating the pH optimum of the enzyme, which was identified to be pH 7.89. At least 90 % of the maximum specific activity could be retained in a range from pH 7.39 to 8.35 (see Figure 3a). A comparison to other IDHs, listed in the BRENDA database reveals this feature to be quite common, as it is close to the median value of pH 8 (see Figure 3b).

**Figure 3.**
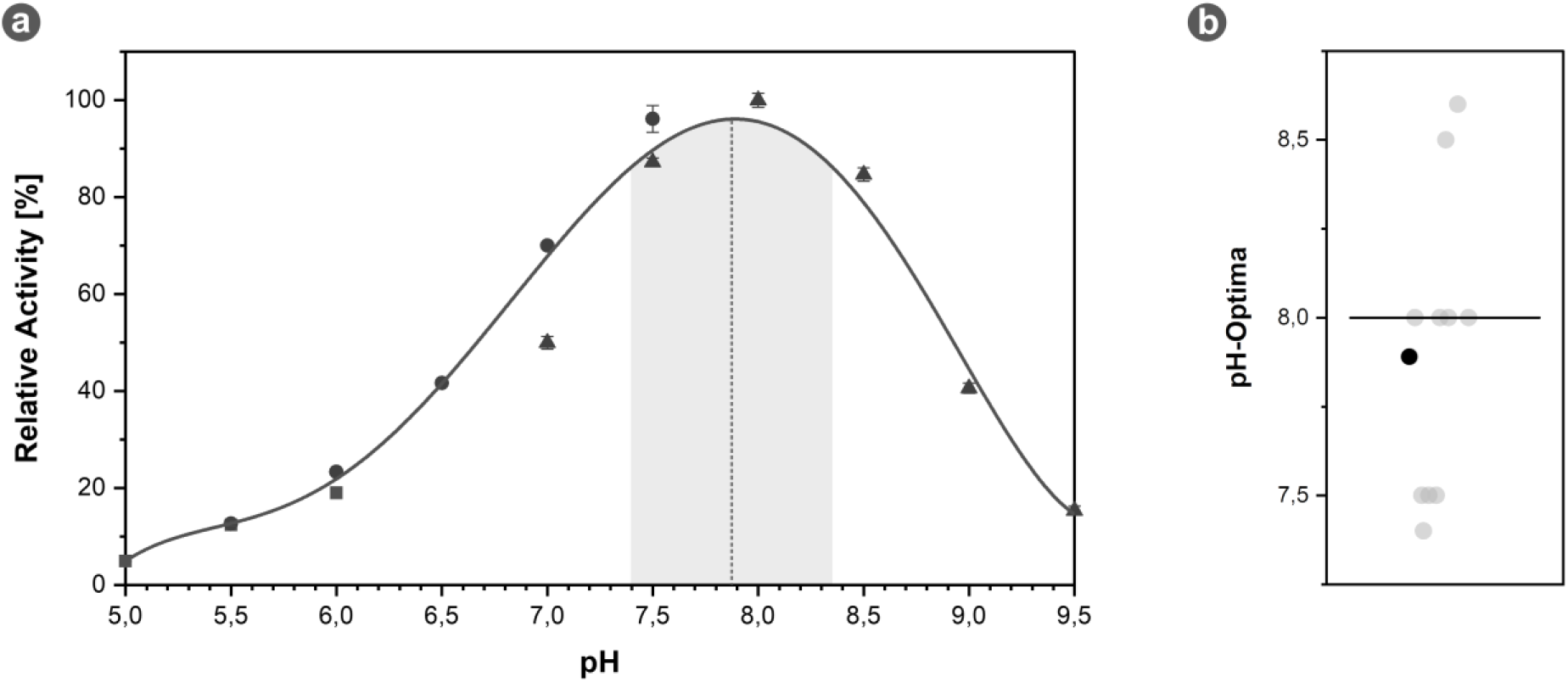
Optimal pH of *Mh*IDH_6x His_. (**a**) Specific IDH activity as a function of the pH value with a polynomial fitting curve of 5th order (R^2^ > 0.99). The global maximum of the curve corresponding to the pH optimum of 7.89 is indicated by a dashed line, the range of specific activity higher than 90 % of the maximal activity is highlighted in grey. pH ranges with sodium acetate (▪), sodium phosphate (•) and TRIS/HCl (▲) buffers are indicated by the respective symbols. Assays were conducted at 28 °C in presence of NADP^+^ and Mg^2+^. (**b**) Distribution of pH optima of homologous IDHs listed in the BRENDA database (see Table A2). The pH optimum of *Mh*IDH_6x His_ is highlighted in black. The median is indicated by a black bar.

Note however, that IDHs in this comparison exclusively originate from neutralophilic organisms, since to our knowledge data on pH optima of IDHs from acidophilic organisms is scarce. One of these few cases being *Thermoplasma acidophilum* IDH (*Ta*IDH). Growing optimally in environments with pH values of 1-2, *T. acidophilum* has a pH_i_ value of 5.8 [57]. Contrary to that, *Ta*IDH displays optimal activity at pH 7.5 [58]. Still, *Ta*IDH is reported to retain a third of its maximal specific activity at pH 5.8 [58], which is not the case for *Mh*IDH_6x His_. Moreover, other enzymes of acidophiles are reported to display highest activity at slightly acidic pH values [6,59,60]. This finding implies a higher intracellular pH of A_DKE compared to other acidophiles.

#### 3.3.3 *Mh*IDH_6x His_ is Characterised by Low NADP^+^ Affinity

Kinetic data of *Mh*IDH_6x His_ was obtained for the substrate DL-isocitrate and the cofactor NADP^+^ (see Figure 4a & b). Overall, kinetic properties of *Mh*IDH_6x His_ regarding DL-isocitrate appear to be quite average compared to other IDHs (see Figure 4c and Table A2), as with *K*_*M*_ = 53.03±5.63 µM, *k*_*cat*_ = 38.48±1.62 s^-1^ and *k*_*cat*_/*K*_*M*_ = 725±107.62 mM^-1^ s^-1^ all parameters lie close to the respective median value. Regarding NADP^+^, on the other hand, *Mh*IDH_6x His_ performs significantly worse in comparison to other IDHs (see Figure 4d and Table A2). A *K*_*M*_ of 1.94±0.12 mM is exceptionally high compared to other IDHs being the least specific enzyme in the comparison. Despite a decent turnover rate close to the median value (*k*_*cat*_ = 43.99±1.46 s^-1^), *Mh*IDH_6x His_ ranks among the three IDHs with the lowest catalytic efficiency (*k*_*cat*_/*K*_*M*_ = 22.69±2.15 mM^-1^ s^-1^). All in all, low affinity to NADP^+^ seems to be the bottleneck limiting the overall reaction rate of *Mh*IDH_6x His_ and possibly the metabolic rate of the whole organism, given that IDH is a key enzyme of the tricarboxylic acid cycle, which is the central metabolic pathway in A_DKE [18].

**Figure 4.**
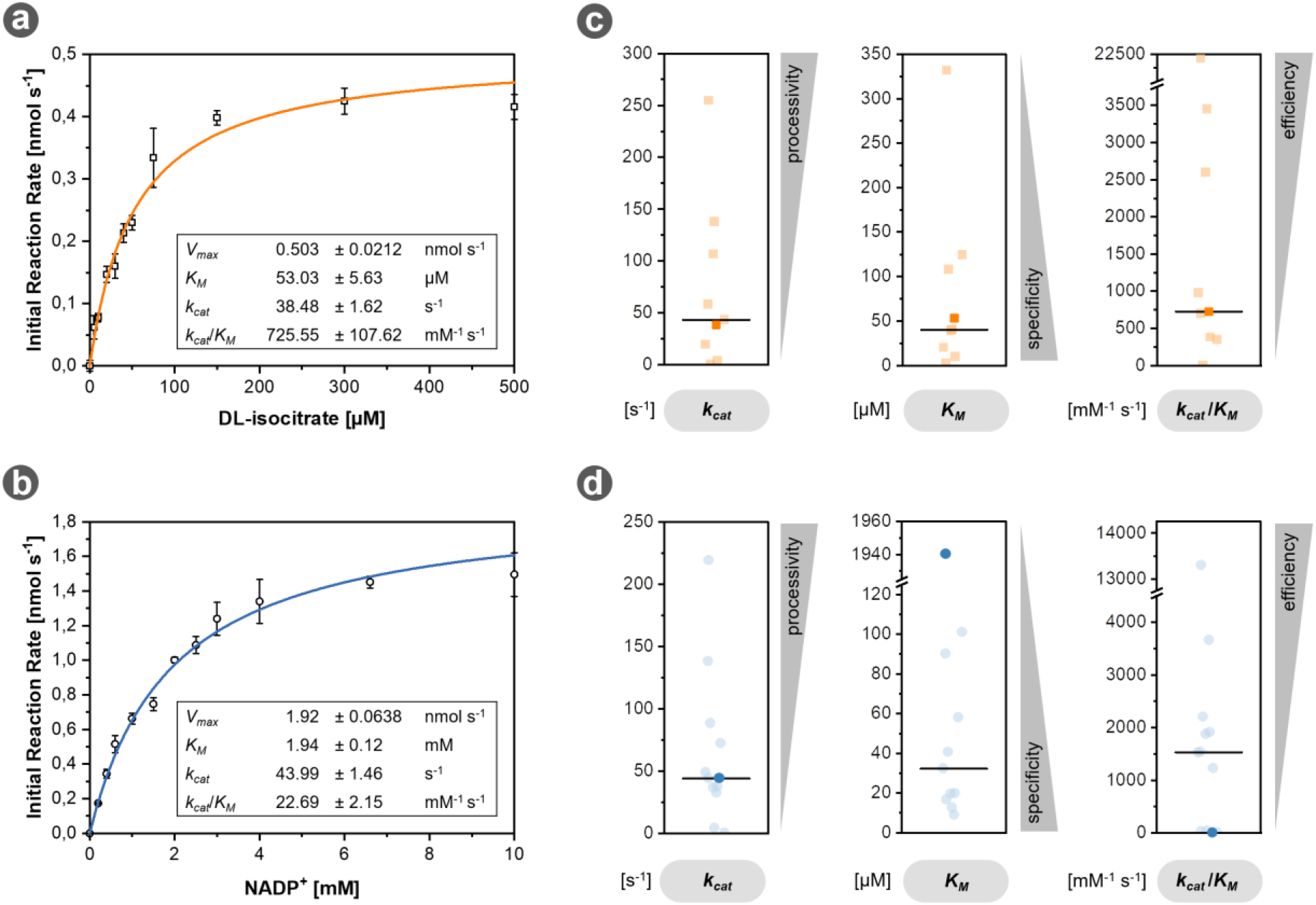
Enzyme kinetics of *Mh*IDH_6x His_. (**a & b**) The initial reaction rate of the enzyme for DL-isocitrate (▫, orange) and NADP^+^ (∘, blue) was measured at the indicated substrate concentrations and fit according to the Michaelis-Menten model (R^2^ (DL-isocitrate) > 0.98, R^2^ (NADP^+^) > 0.99). The corresponding kinetic parameters derived from the fits are given in the respective inset tables. All assays were conducted at 28 °C in presence of Mg^2+^, as well as 1 mM of DL-isocitrate and 20 mM of NADP^+^, respectively. Reaction mixtures contained 2 and 0.6 µg enzyme per reaction for NADP^+^ and DL-isocitrate kinetics, respectively. (**c & d**) Comparison of kinetic parameters of *Mh*IDH_6x His_ for DL-isocitrate (▫, orange) and NADP^+^ (∘, blue) with those of other IDHs listed in the BRENDA database (transparent, see Table A2). The parameters of *Mh*IDH_6x His_ are highlighted in opaque orange and blue, respectively. The corresponding median values are indicated by a black bar.

To investigate potential ligand binding mechanisms in *Mh*IDH, we conducted a multiple sequence alignment with other experimentally verified IDHs and modelled a putative structure (see Figure 5) using a crystal structure of *Ec*IDH as a template. The model features high estimated local model quality and shows the characteristic fold of prokaryotic NADP-dependent IDHs, comprising a large and a small domain responsible for cofactor and substrate binding, respectively, as well as a clasp domain allow-ing homo-dimerisation ([52,53], see Figure 5b). The estimated local B-factor of the model indicates a rigid core, as well as flexible loops surrounding the active site in between the small and large domain (see Figure 5b), which allow conformational change necessary for catalytic activity in *Ec*IDH [32]. Ligands isocitrate, NADP^+^ and Mn^2+^ could be docked in the active sites of the homo-dimeric model, with their relative positions closely resembling those in *Ec*IDH (see Figure 5c).

**Figure 5.**
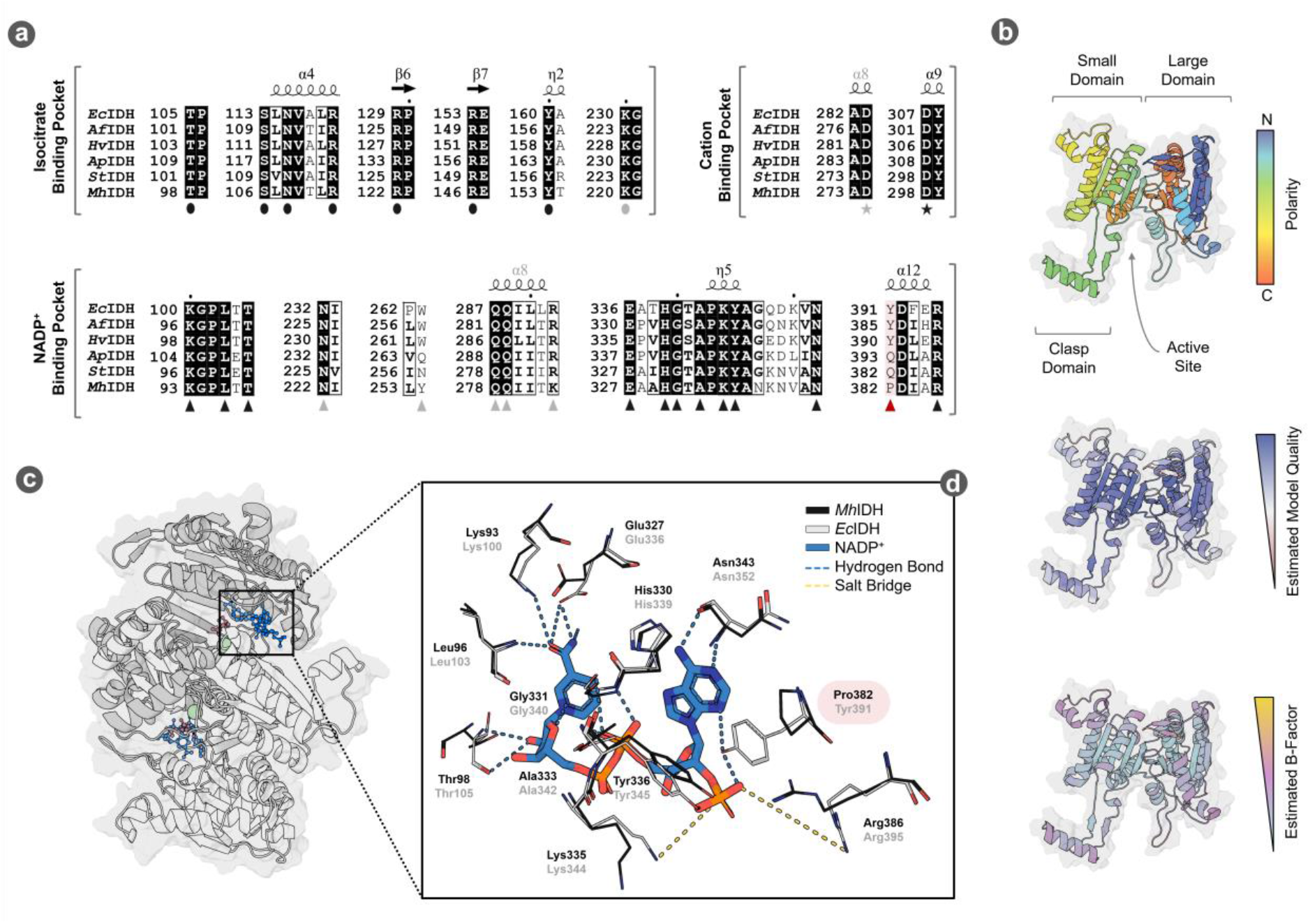
Putative structure and ligand binding in *Mh*IDH. (**a**) Partial multiple sequence alignment of the substrate and cofactor binding pockets of *Mh*IDH with IDH sequences from *E. coli* K-12 (*Ec*IDH, NCBI: P08200. 1), *Archaeoglobus fulgidus* DSM 4304 (*Af*IDH, NCBI: O29610.1), *Haloferax volcanii* DS2 (*Hv*IDH, NCBI: D4GU92.1), *Aeropyrum pernix* K1 (*Ap*IDH, NCBI: GBF08417.1) and *Sulfolobus tokodaii* strain 7 (*St*IDH, NCBI: BAB67271.1). Identical amino acids are highlighted in black, homologous amino acids are boxed. Residues involved in isocitrate (•) cation (⋆) and NADP^+^ (▴) binding in *Ec*IDH according to [32] are highlighted by the corresponding symbols. The position of Pro382 in *Mh*IDH is highlighted in red. Residues of the second homo-dimer subunit involved in ligand binding are highlighted in grey symbols. Full alignment see Figure A1. (**b**) Putative structure of monomeric *Mh*IDH homology-modelled after the crystal structure of *Ec*IDH ([32], PDB: 4AJ3, 49.5 % sequence homology, 1.9 Å resolution) in ribbon representation and coloured according to orientation of the backbone, as well as estimated local model quality and B-factor as determined by the ResQ server. The surface representation of the protein is indicated in the background. (**c**) Ribbon representation of a putative quaternary structure of *Mh*IDH in top view, forming a homo-dimer with an active site located between the large and small domain of each subunit. Docked ligands isocitrate (red), NADP^+^ (blue) and Mn^2+^ (green) are shown in ball-and-stick representation. (**d**) Detail-view of a structural alignment of the NADP binding pockets in the *Mh*IDH model (black) and the *Ec*IDH crystal structure (grey). Side chains of amino acids presumably involved in cofactor binding, as well as NADP^+^ are displayed as stick-models and are highlighted according to their atomic composition: O – red; N – blue, P – orange; C – grey (*Ec*IDH), black (*Mh*IDH) or blue (NADP^+^). Interactions between *Ec*IDH residues and NADP^+^ are indicated by dashed lines (salt bridges – yellow; hydrogen bonds – light blue).

A comparably average *K*_*M*_ value for isocitrate is not surprising, considering that without exception all amino acids known to be involved in isocitrate binding in other IDHs [32,52,53,56] are conserved in the isocitrate binding pocket of *Mh*IDH (see Figure 5a).

Furthermore, low affinity of *Mh*IDH for NADP^+^ can be explained by structural analysis, as well. The NADP^+^ binding pocket in *Ec*IDH is formed by the 3_10_-helix η4 (residues 318-324), the NADP^+^ binding loop (residues 336-352), as well as helix α12 (residues 390-397) [32]. In particular, amino acids Lys100, Leu103, Thr105, Asn232*, amino acids 258*-261*, Trp263*, Gln287*, Gln288*, Arg292*, Glu336, His339, Gly340, Ala342, Lys344, Tyr345, Asn352, Tyr391 und Arg395 (* marks amino acids from the second subunit of the homo-dimer) are involved in binding NADP^+^ via hydrogen bonds or salt bridges ([32], see Figure 5a). Corresponding residues in *St*IDH [52] and *Ap*IDH [53] have been described to facilitate NADP^+^ binding, as well (see Figure 5a). Almost all of the corresponding amino acids in *Mh*IDH are conserved or at least dis-play similar physicochemical properties (Tyr254* instead of Trp263* and Lys282* in-stead of Arg292*), the only exception being Tyr391 (see Figure 5a & d), which is substituted for a proline in MhIDH (Pro382). While this appears to be a common feature among isopropylmalate dehydrogenases rather than IDHs (i.e. in *Thermus thermophilus* [61]), *Mh*IDH showed significantly higher sequence homology to the latter (see Table A1). Since Tyr391 forms hydrogen bonds stabilising the 2’-phosphate of NADP^+^ (see Figure 5a & d) this amino acid plays a critical role in cofactor stabilisation and selectivity in *Ec*IDH [50,51,61]. Moreover, it has been reported that a proline at this position disrupts the local α-helix in favour of a β-turn [61,62], which could distance Lys386, another crucial residue ensuring NADP^+^ specificity, from the 2’-phosphate of NADP^+^ and thereby decrease cofactor stabilisation even more.

## 4. Conclusion

Although several approaches lead to new findings about Micrarchaeota in the last decade, the survival strategies of these ultra-small, acidophilic organisms are still not fully understood. In this study, we gained evidence for the internal pH of “Ca. Micrarchaeum harzensis A_DKE”, by characterisation of its IDH. The enzyme was successfully produced in *E. coli* and biochemically characterised. Compared to other known IDHs, the NADP^+^ and divalent cation-dependent protein from A_DKE seems to be highly inefficient because of the amino acid composition of its NADP^+^ binding-pocket. Since *Mh*IDH plays a role in A_DKE’s main pathway for generation of reducing equivalents, its inefficiency is in line with the slow growth rates of the Micrarchaeon.

Over the years, a vast arsenal of methods, suitable for the determination of pH_i_ values has been developed, including cell homogenate measurement, pH-sensitive fluorescent proteins and fluorescent probes, injection of microelectrodes and ^31^P-NMR spectroscopy. All these methods come with individual strengths and weaknesses, discussed elsewhere [11,63]. In our specific case, however, experimental determination of the pH optimum of an intracellular enzyme as described in [6,43] remained the only viable option.

The presented data suggests that A_DKE maintains a slightly alkaline cytosolic milieu close to pH 8, while thriving in acidic environments of pH 2, resulting in a steep pH gradient of several orders of magnitude. Should this assumption be correct, A_DKE would have the highest pH_i_ among all acidophiles described so far, which raises the question how the Micrarchaeon is able to maintain this pH gradient. In literature there are several synergistic strategies of proton homoeostasis described for acidophiles [5,11], many of which might apply to A_DKE as well:

Membranes consisting of archaeal tetraetherlipids have been reported to be highly impermeable for protons [5,64–67]. With a caldarchaeol content of 97 % the cell membrane of A_DKE is predominantly composed of such lipids [17]. Furthermore, in *T. acidophilum* HO-62 a correlation between acid tolerance and elevated levels of surface glycosylation has been found [68]. The cell surface of A_DKE is mostly covered by a proteinaceous, heavily glycosylated S-layer [21]. Since, biomimetic experiments strongly suggest, that polysaccharide chains attached to the cell surface might effectively create a proton shelter [69], the glycans linked to A_DKE’s S-layer, could allow further shielding from protons. Another mechanism of acidophiles to repel invading protons is the formation of a positive potential at the inside of the cell membrane via cation transporters [5,11,12,67]. Also, acidophiles are known to express a variety of primary proton transporters in order to counteract cellular protonation caused by ATP synthase activity [5,11,12]. According to transcriptomic data, A_DKE seems to express several genes encoding (putative) proton pumps and cation transporters (see Table A3), which would allow export of protons to the extracellular space, as well as antiport exchanging protons for cations.

Lastly, this study proves the viability of recombinant production of functional A_DKE proteins in *E. coli*, which opens numerous possibilities for the biochemical characterisation of proteins of unknown function in A_DKE.

## Author Contributions

Conceptualisation, J.G.; methodology, D.W.; validation, D.W. and J.G.; formal analysis, D.W.; investigation, D.W.; resources, J.G.; data curation, D.W. and S.G.; writing—original draft preparation, D.W., S.G. and J.G.; writing—review and editing, D.W., S.G. and J.G.; visualisation, D.W.; supervision, J.G.; project administration, J.G.; funding acquisition, J.G. All authors have read and agreed to the published version of the manuscript.

## Funding

This research received no external funding.

## Data Availability Statement

All data shown is contained within the article.

## Conflicts of Interest

The authors declare no conflict of interest.

## Appendix A

**Figure A1.**
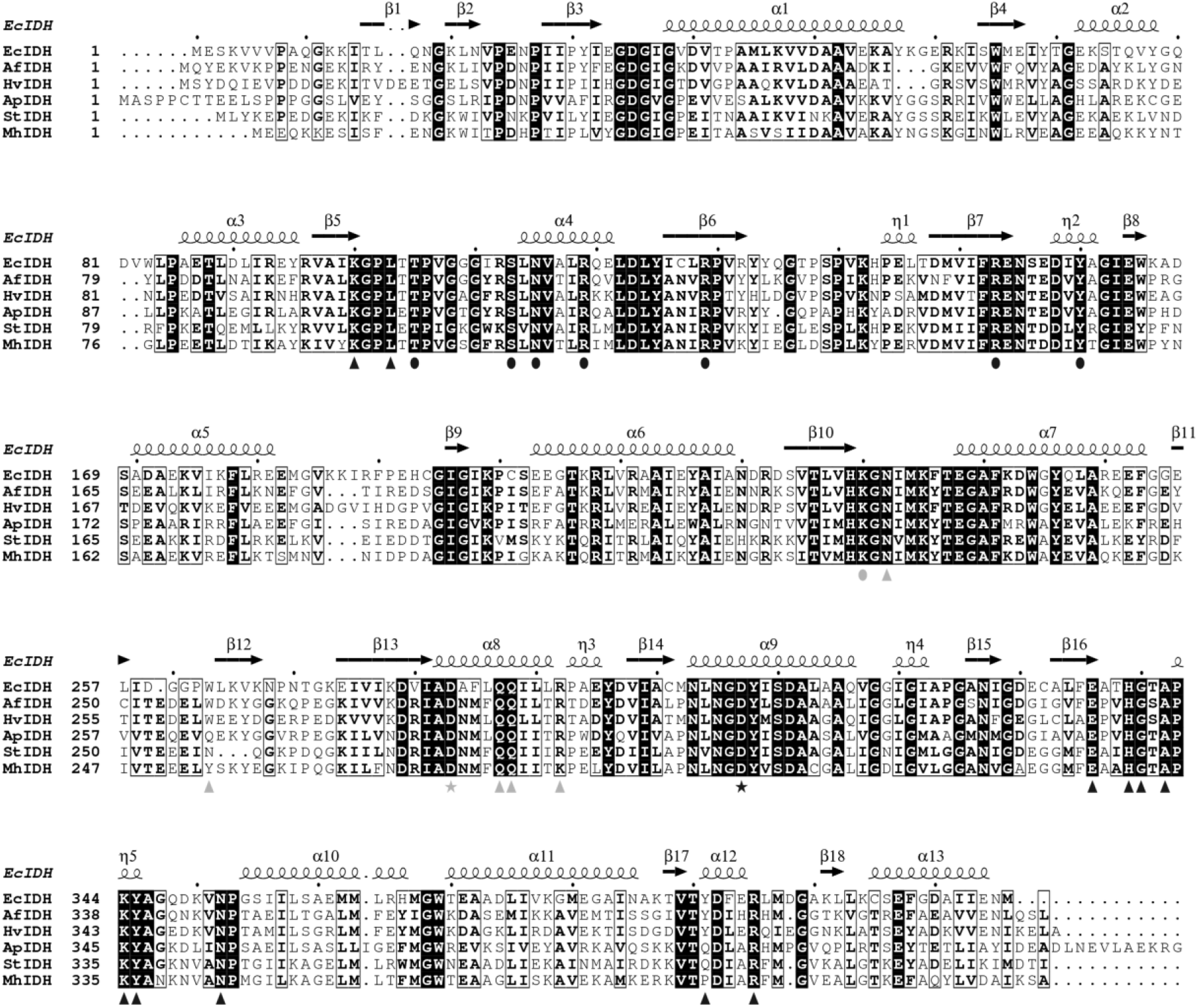
Multiple Sequence Alignment of *Mh*IDH with homologous NADP-specific IDHs. *Mh*IDH-homologues from *E. coli* K-12 (*Ec*IDH, NCBI: P08200.1), *Archaeoglobus fulgidus* DSM 4304 (*Af*IDH, NCBI: O29610.1), *Haloferax volcanii* DS2 (*Hv*IDH, NCBI: D4GU92.1), *Aeropyrum pernix* K1 (*Ap*IDH, NCBI: GBF08417.1) and *Sulfolobus tokodaii* Strain 7 (*St*IDH, NCBI: BAB67271.1) were identified via BLASTp-search and aligned using Clustal Omega. Identical amino acids are highlighted in black, similar amino acids are boxed. Secondary structure elements of *Ec*IDH (above) named according to their type and number of appearance are indicated by arrows (β-strands), as well as large and small squiggles (α- and 3_10_ (η)-helices), respectively. Amino acids involved in isocitrate-(•), NADP^+^- (▴) and cation-binding (⋆) in *Ec*IDH are highlighted below by the indicated symbols. The symbols in grey represent amino acids which interact with the ligands in the active site of the other homo-dimer subunit.

**Table A1.**
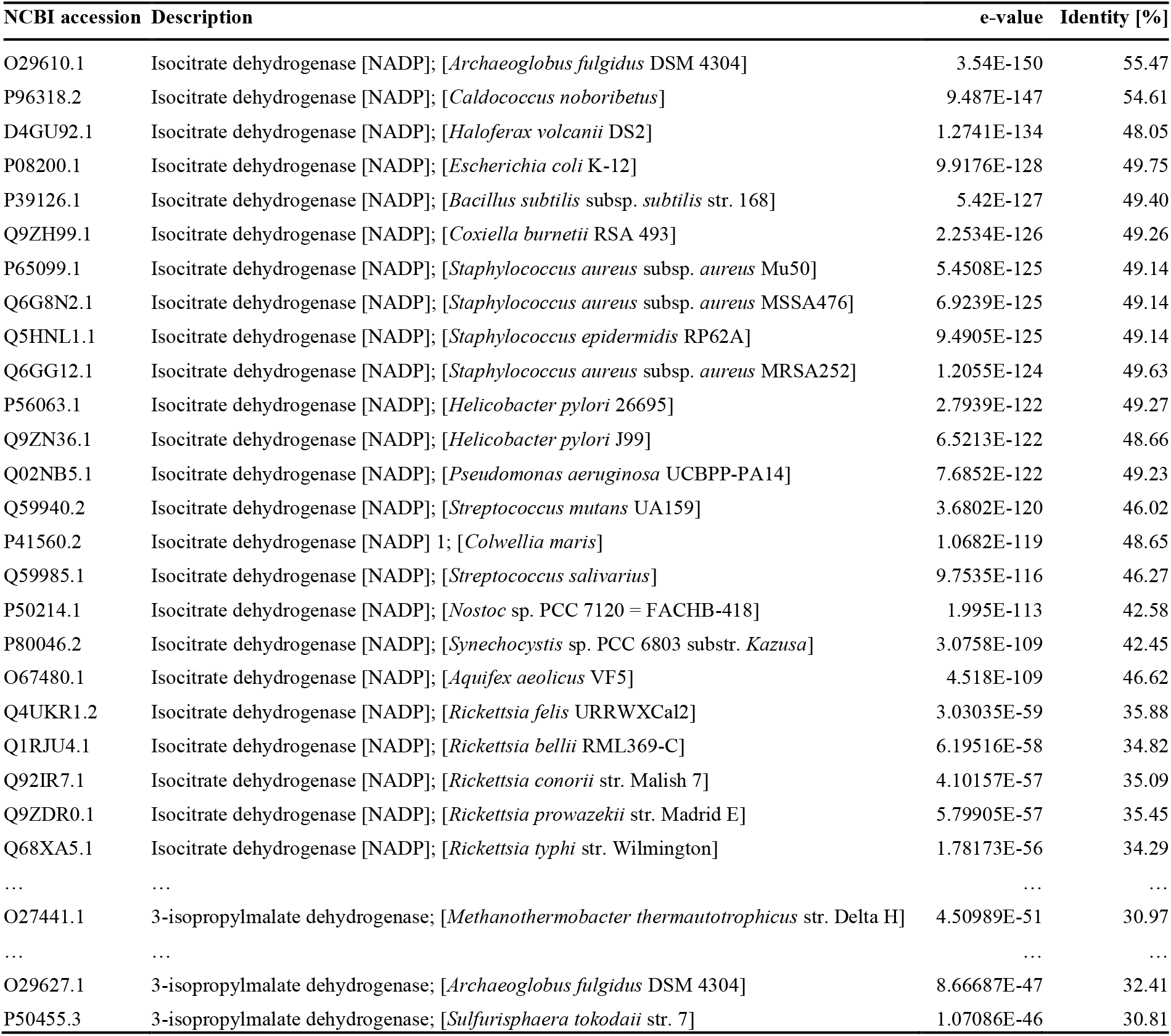
Selected results of a BLASTp-search for homologues of *Mh*IDH using the UniprotKB/swiss-prot database as a reference.

**Table A2.**
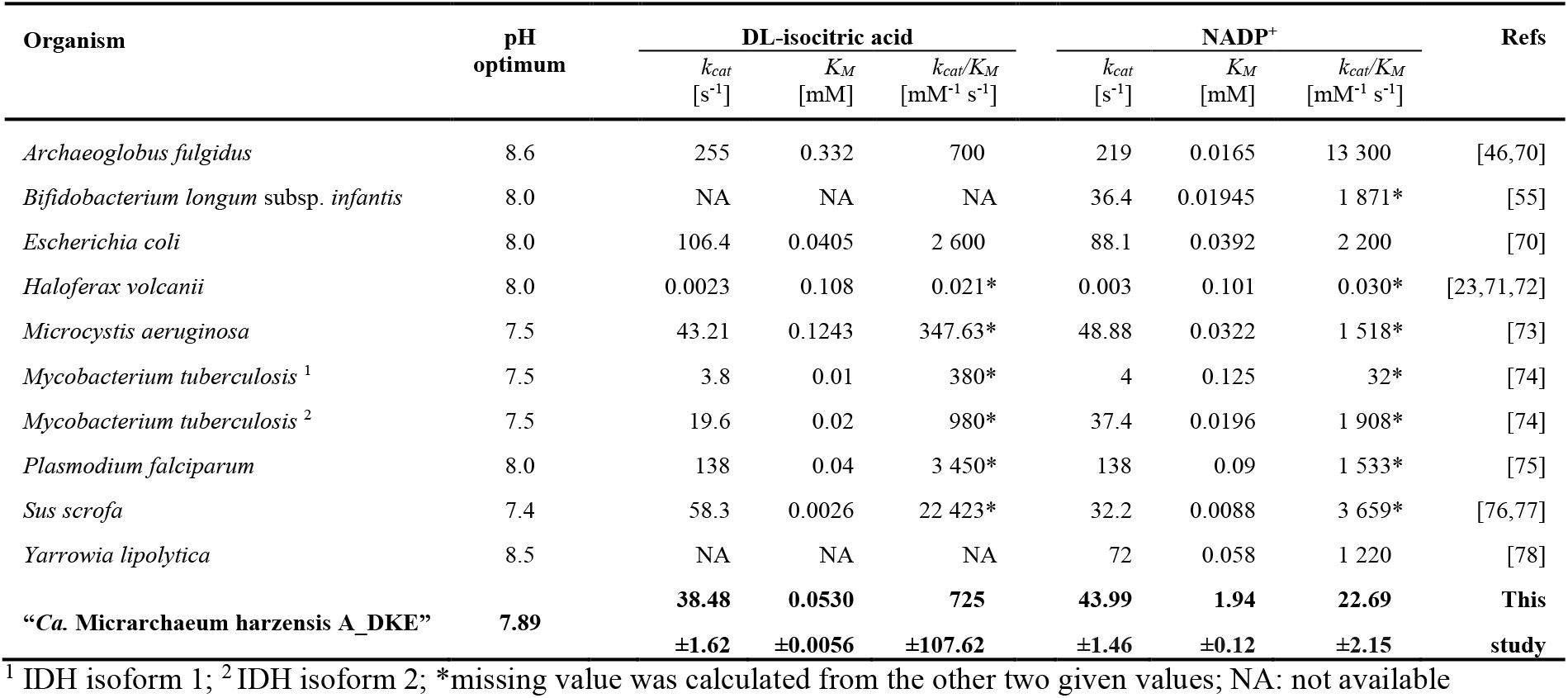
Overview on NADP-dependent IDHs with a mostly complete set of catalytic parameters listed in the BRENDA database.

**Table A3.** List of known and putative H^+^- and cation-transporters encoded in the A_DKE genome. Given with the respective e-values for KEGG Orthology (KO) prediction and transcriptomic TPM-values indicating their relative expression levels. Data taken from [17].

| Gene | Putative function | KO identifier | e-value | TPM |
| --- | --- | --- | --- | --- |
| Micr_00103 | manganese transport protein | K03322 | 7.6E-29 | 0.31 |
| Micr_00140 | sodium transport system permease protein | K09696 | 4.2E-21 | 0.24 |
| Micr_00144 | MscS family membrane protein | K16052 | 1.8E-30 | 0.77 |
| Micr_00155 | voltage-gated potassium channel | K10716 | 1.2E-28 | 0.71 |
| Micr_00226 | inorganic pyrophosphatase | K01507 | 1.00E-38 | 4.22 |
| Micr_00278 | ammonium transporter, Amt family | K03320 | 7.6E-43 | 0.06 |
| Micr_00279 | ammonium transporter, Amt family | K03320 | 4.9E-21 | 0.05 |
| Micr_00330 | MFS transporter, PHS family, inorganic phosphate transporter | K08176 | 5.3E-121 | 0.52 |
| Micr_00498 | inorganic pyrophosphatase | K01507 | 5.9E-19 | 0.42 |
| Micr_00551 | vacuolar iron transporter family protein | K22736 | 5.4E-23 | 0.86 |

## Notes

### Competing Interest Statement

The authors have declared no competing interest.

